# Genetic diversity, evolutionary dynamics and pathogenicity of ferret badger rabies virus variants in mainland China, 2008-2018

**DOI:** 10.1101/2021.04.19.440555

**Authors:** Faming Miao, Jinghui Zhao, Nan Li, Ye Liu, Teng Chen, Lijuan Mi, Jinjin Yang, Qi Chen, Fei Zhang, Jie Feng, Shunfei Li, Shoufeng Zhang, Rongliang Hu

## Abstract

In contrast to dog associated human rabies cases decline year by year due to the rabies vaccination coverage rates increase in China, ferret badger (FB, *Melogale moschata*)–associated human rabies cases emerged in the 1990s, and are now an increasingly recognized problem in southeast China. To investigate epidemiology, temporal evolution dynamics, transmission characterization and pathogenicity of FB-associated rabies viruses (RABVs), from 2008 to 2018, we collected 3,622 FB brain samples in Jiangxi and Zhejiang Province, and detected 112 RABV isolates. Four FB-related lineages were identified by phylogenetic analysis (lineages A–D), the estimated Times to Most Recent Common Ancestor were 1941, 1990, 1937 and 1997 for lineages A–D respectively. Furthermore, although no FB-associated human rabies case has been reported there apart from Wuyuan area, FB-RABV isolates are mainly distributed in Jiangxi Province. Pathogenicity of FB-RABVs was assessed using peripheral inoculation in mice and in beagles with masseter muscles, mortality-rates ranging from 20% to 100% in mice and 0 to 20% in beagles in the groups infected with the various isolates. Screening of sera from humans with FB bites and no postexposure prophylaxis to rabies, revealed that 5 of 9 were positive for neutralizing antibodies of RABV. All the results above indicated that FB-RABV variants caused a lesser pathogenicity in mice, beagles and even humans. Vaccination in mice suggests that inactivated vaccine or recombinant subunit vaccine products can be used to control FB- associated rabies, however, oral vaccines for stray dogs and wildlife need to be developed and licensed in China urgently.

**IMPORTANCE:** In recent years FB-associated rabies virus has been identified as a major life-threatening pathogen in some districts in China. To understand the risk to public health and the contemporary dynamics, the present study conducted extensive investigations on FB rabies, distribution, virus isolation, phylogeny analysis, pathogenicity determination of various FB rabies virus strains, besides, serologic epidemiology survey to those whom bit by FB was also collected. The results show that the majority of FBs dwell in southeast China, like Jiangxi and Zhejiang Province, Phylogenetic analysis indicates that all isolated FB RABVs evolved from dogs, and the FB RABV can separate into 4 distinct lineages distributed relatively independent in different areas. The isolate strains differ in pathogenicity, although they have relatively lower pathogenicity compared to dog rabies virus according to our study, the need for further study to licensed oral vaccines and FB RABV pathogenesis is emphasized in order to control rabies.

---

Rabies is an acute, progressive and encephalitic disease. Once clinical signs manifests, the disease usually is considered to be invariably fatal. It continues to present public health problems worldwide and causes more than 60,000 human deaths annually (1). The majority of these deaths are in developing countries, primarily in Africa and Asia, more than 95% of human rabies cases are attributed to dog bites worldwide and the ration is higher in developing countries (2). However, in southeast China, the percentage dog-associated human rabies is relatively low, up to 80% of the reported human rabies cases were inferred to be caused by ferret badger (FB) bites in some districts in Zhejiang and Jiangxi province (3, 4).

The Chinese ferret badger, which dwells mainly in southeastern China. These mustelids have several names like crab-eating mongoose, rice field dog, viviparid-eating dog, loach-eating dog, and white face weasel—mainly because of their omnivorous behavior and external appearance. FB-associated human rabies cases in China were emerged in 1994 (5), forty-six human rabies cases exposed to FB bites were reported in Zhejiang and Jiangxi province from 1994-2007 (6).

In 2008, we isolated the first FB rabies virus strain, since then, we continued our surveillance, and collected additional positive samples mainly distributed in Jiangxi and some in Zhejiang province that were isolated from FB. We have conducted the laboratory-based surveillance, virus isolation, nucleotide sequencing and molecular characterization analysis (7–9). Recently, we investigated the temporal dynamics of FB-RABVs in China based on N sequences from samples collected from 2014 to 2016 (10), and demonstrated that the increasing genetic diversity of RABVs in FBs is a result of the selective pressure from coexisting dog rabies virus, and FB rabies is likely occurring as an independent enzootic that became established in the FB populations from a dog RABV variant distributed in southeast China (11), which served as a reservoir for human infections and a source of new variants capable of emerging or reemerging in the future; however, little is known about the genetic diversity of FB-associated RABVs, their evolutionary dynamics and the role of FBs in maintaining RABV transmission in the epizootic regions, especially the pathogenicity of FB RABV remains to be unclear.

In this study, we collected 3,622 FB brain samples from 2008 through 2018 in these regions and detected 112 FB-RABV isolates from the collected samples, and sequenced the full-length of the nucleoprotein (N) gene from each isolate, constructed the phylogenetic and temporal evolution dynamics analysis. Furthmore, we determined the pathogenic characteristics of FB RABV in mice and beagles, tested the serum antibody titers for RABV who bit by FBs. Moreover, we reported that using inactivated vaccines or adenovirus recombinant vaccine can protect mice from lethal challenge with FB RABV infection.

## Results

### FB RABVs were constantly detected over the years in Poyang and Qiandao lake regions

As depicted in Table 1, from 2008 to 2018, FB-associated RABV nucleic acids were regularly detected in our collected samples from the Poyang lake region, with a positive rate ranging from 1.7%–6.3%. In the Qiandao lake region, we detected 5 RABV positive among 138 samples from 2008 through 2013 (Table 1). A total of 112 FB RABV strains were detected and 70% infectious viruses were successfully isolated from all positive tissues and cultured in suckling mouse brain. The full-length of the N gene from each positive sample was amplified by RT-PCR and sequenced for phylogenetic and evolutionary studies.

**Table 1.** Rabies virus detection in ferret badgers in Poyang and Qiandao lake regions, 2008-2018.

| Year | Positive numbers/Total tested numbers (% positive) |  |  |
| --- | --- | --- | --- |
|  | Poyang Lake | Qiandao Lake | Total |
| 2008 | 4/63 (6.3) | 1/6 (16.7) | 5/69 (7.2) |
| 2009 | 10/294 (3.4) | 0/63 | 10/357 (2.8) |
| 2010 | 4/148 (2.7) | 0/9 | 4/157 (2.5) |
| 2011 | 6/248(2.4) | 0/20 | 6/268 (2.2) |
| 2012 | 13/751 (1.7) | 1/19 (5.2) | 14/770 (1.8) |
| 2013 | 24/479 (5.0) | 3/21 (14.2) | 27/500 (5.4) |
| 2014 | 4/126 (3.2) | / | 4/126 (3.2) |
| 2015 | 1/36 (2.8) | / | 1/36 (2.8) |
| 2016 | 10/363 (2.8) | / | 10/363 (2.8) |
| 2017 | 24/677 (3.5) | / | 24/677 (3.5) |
| 2018 | 7/299 (2.3) | / | 7/299 (2.3) |
| Total | 107/3484 (3.1) | 5/138 (3.6) | 112/3622 (3.1) |

### Segregation of FB RABVs into geographic clades and evidence of RABV cross species transmission between FBs and dogs

The 112 FB-associated RABVs were classified into 4 lineages (A–D) using data from 132 Chinese RABV N gene sequences, and each lineage was grouped geographically by the location of the virus isolate (Fig 1). Lineage A, with 41 FB-associated RABV isolates, 40 distributed in east of Poyang lake and 1 distributed in Zhejiang province, was embedded in China canine RABV Group II (China II). The FB RABVs in lineage A were closely related to Chinese vaccine strains CTN181 and CTN7, which were derived from a dog street virus in the 1950s (12).

**Fig 1.**
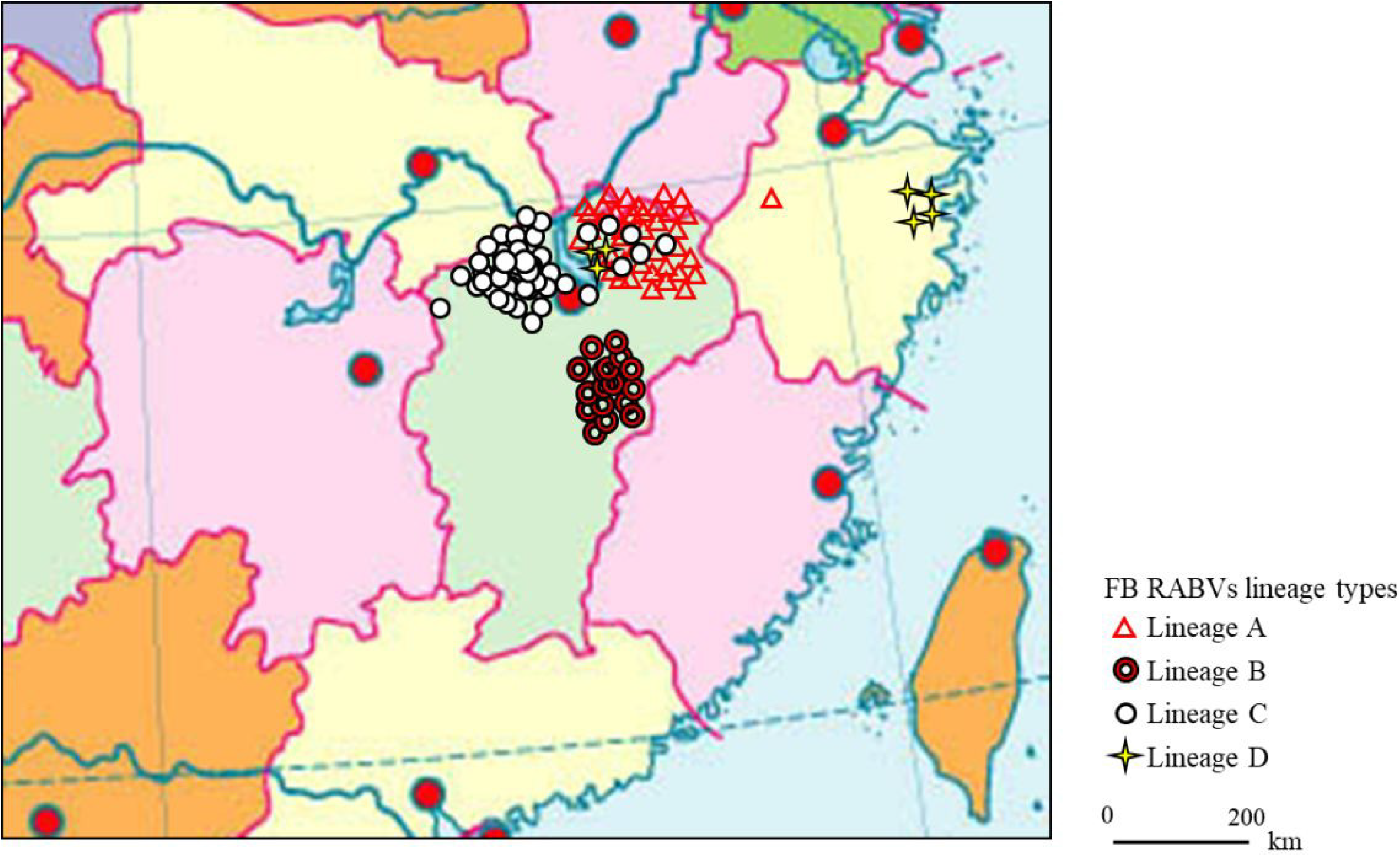
Continental distribution of ferret badger rabies virus isolates in southeast China during 2008-2018. The 41 FB rabies virus isolates in Lineage A were mainly distributed in Qiandao lake (Zhejiang province) and east Poyang lake (Jiangxi province) regions. Lineage B had 17 isolates from Fuzhou district, south of Poyang lake. Lineage C included 47 isolates, 41 distributed in west Poyang lake, and the other 6 in east Poyang lake, mixing geographically with Lineage A. Lineage D had 4 isolates in Taizhou district, Zhejiang province and 3 in east Poyang lake.

Lineage B, C and D were grouped with China canine RABV Group I (China I) (13). Lineage B had 17 isolates from the Fuzhou district, south of Poyang lake (7). Lineage D had 7 FB isolates, 4 FB isolates (ZJ12-03, ZJ13-66, ZJ13-130, ZJ13-431), located in Taizhou district, Zhejiang province, other 3 isolates located in east Poyang lake. Of note, a canine RABV isolate denoted as LH from Zhejiang province, where Qiandao lake is located, was embedded in the FB lineage D, and lineage D and canine RABV Group closely related phylogenetically. Lineage C included 47 FB-associated RABVs, from 2008-2018, 41 isolated from west Poyang lake, and the other 6 from east Poyang lake, mixing geographically with Lineage A (Fig 1). However, lineages C and A were phylogenetically distinctly separate. The 47 isolates within Lineage C, the majority lacated in the west of Poyang lake, shared approximately 95% nucleotide identity with China canine RABV Group I, and 88% nucleotide identity with China canine RABV Group II.

### The time of FB RABV emergence in China by evolutionary analysis

The Bayesian MCMC approach used assumed a constant population size and an uncorrelated log normal molecular clock. The mean rate of nucleotide substitution for the N gene was 5.19×10^−4^ substitutions per site per year (95% HPD values, 3.48– 7.05×10^−4^) which is close to that reported previously (14, 15). The calculated the Time of the Most Recent Common Ancestor (TMRCA) of China RABVs was 1745 CE (95% HPD values, 1700–1863). For FB Lineage A, the TMRCA was 1941 (95% HPD 41– 100 years, or year 1910–1969). For Lineage B, the TMRCA was 1990 (95% HPD 12–28 years, or year 1982–1998). For Lineage C, the TMRCA was 1937 (95% HPD 42–104 years, or year 1906–1968). For Lineage D, the TMRCA was 1997 (95% HPD 9– 18 years, or year 1992–2001). Lineages B and D could be recent events diverged from dog RABV spillovers to the FB populations, while lineages A and C could represent old historical spillovers with subsequent establishment in the FB population. In our 3,622 FB samples, the chance of detecting a FB RABV isolate in Lineages A and C was 3.7 fold higher than finding a RABV in Lineages B and D (88 versus 24 in the 3622 samples).

### Pathogenicity of FB-RABV isolates in mice and beagles

Following intramuscular (IM) inoculation of 5 week-old mice, all mice infected with the JX08-45, JX13-189 and BD06 strains died within 6-11 dpi after showing typical neurological symptoms like the appearance of tremble, shaking, anger, then later hind limb paralysis of one or both limbs. However, the severities of infection were clearly different in other isolates (Table 2). The weight loss occurred during the disease progression from the onset of clinical signs to the point of death. Infected mice lost on average 19.3-32% of their overall body weight during the disease progression (data not shown). The incubation periods of the isolates varied from 4 to 12 days: JX13-189 had the shortest incubation period; mice infected with JX13-417 and ZJ12-03 had nearly the same incubation periods and clinical signs; and JX13-228 and JX10-37 infected mice had the longest incubation period (Table 2).

**Table 2.** Morbidity, incubation and morbidity period of mice following intramuscular inoculation with each strain.

| Strain | Lineage | MICLD<sub>50</sub><br>log10 | Fatality/<br>inoculated | Incubation<br>period (days) |  | Mean morbidity<br>period (days) |
| --- | --- | --- | --- | --- | --- | --- |
|  |  |  |  | r | mean |  |
| BD06 | China I | 5.2 | 5/5 | 6-8 | 6.6 | 1.2 |
| JX13-189 | FB-Lineage A | 4.9 | 5/5 | 3-5 | 4 | 2 |
| JX08-45 | FB-Lineage A | 4.9 | 5/5 | 4-7 | 6 | 2.6 |
| JX13-417 | FB-Lineage B | 4.5 | 3/5 | 7-13 | 10.7 | 2.3 |
| JX09-17 | FB-Lineage B | 4.4 | 3/5 | 5-8 | 6.7 | 3.7 |
| JX13-228 | FB-Lineage C | 4.4 | 1/5 | 12 | 12 | 6 |
| JX10-37 | FB-Lineage C | 4.6 | 3/5 | 11-13 | 12 | 6 |
| ZJ12-03 | FB-Lineage D | 4.4 | 4/5 | 7-11 | 9.8 | 2.2 |
| ZJ13-431 | FB-Lineage D | 4.6 | 4/5 | 5-6 | 5.8 | 2 |

To further evaluate their pathogenicity in dogs, four groups of female beagles (n = 5) were inoculated intramuscularly into masseter muscles, all animals were observed for 90 days post-inoculation. The mortality-rate of Zhejiang isolates, ZJ13-431, was 20% (1/5). Other 2 FB-RABV isolates, JX13-189 and JX13-228, exhibited no pathogenicity in beagles. Six of fifteen survived animals had detectable virus neutralizing antibodies (VNA) for rabies (0.13-2.6 IU/mL). After challenge with lethal BD06, rabies VNA of all the surviving animals had seroconverted (0.29-53.3 IU/mL) (Table 3).

**Table 3.** Rabies virus neutralizing antibody titers (IU/mL) and mortality of beagles after infection.

| Group | Fatality/Total | Incubation<br>period (days) | VNAs (IU/mL) at days p.i. |  | After challenge |
| --- | --- | --- | --- | --- | --- |
|  |  |  | 0 | 30 | 14 days |
| BD06 | 4/5 | 9-16 | 0 | 0 | 53.3 |
| ZJ13-431 | 1/5 | 19 | 0 | 0 | 1.5 |
|  |  |  | 0 | 0 | 13.5 |
|  |  |  | 0 | 0.29 | 40.5 |
|  |  |  | 0 | 0 | 4.5 |
| JX13-189 | 0/5 | — | 0 | 0.29 | 1.97 |
|  |  |  | 0 | 0 | 0.87 |
|  |  |  | 0 | 0.13 | 1.5 |
|  |  |  | 0 | 0 | 1.14 |
|  |  |  | 0 | 2.6 | 5.92 |
| JX13-228 | 0/5 | — | 0 | 0 | 0.5 |
|  |  |  | 0 | 0 | 40.5 |
|  |  |  | 0 | 2.6 | 2.6 |
|  |  |  | 0 | 0 | 0.29 |
|  |  |  | 0 | 0.29 | 4.5 |

### RABV neutralizing antibodies was detectable in people with frequent contacts of FBs

The nine human volunteers were selected from persons with frequent contacts with FB of more than 3 years’ duration. Testing of their sera revealed that 3 of the 9 had high titers of rabies virus–neutralizing antibodies (>2.6IU/ml), and another two persons generated a certain amount of VNA titers (Table 4).

**Table 4.** Results of serologic rabies virus neutralizing antibody titers (IU/mL) in 9 volunteers from Jiangxi province.

| Volunteer No. | Area | Sex | Age yr | Time of bites | Rabies virus neutralizing antibodies (IU/mL) |
| --- | --- | --- | --- | --- | --- |
| H1 | Fengxin County, Nanchang | M | 47 | multiple | 2.6 |
| H2 | Gaoan County, Jiujiang | F | 47 | multiple | 0 |
| H3 | Anyi County, Nanchang | M | 40 | multiple | 17.77 |
| H4 | Anyi County, Nanchang | M | 62 | multiple | 0.29 |
| H5 | Yongxiu County, Yichun | M | 50 | multiple | 0 |
| H6 | Nanchang | M | 26 | multiple | 10.26 |
| H7 | Hukou County, Jiujiang | M | 46 | multiple | 0.17 |
| H8 | Nanchang | M | 47 | twice | 0 |
| H9 | Jinxian County, Nanchang | M | 60 | multiple | 0 |

### Rabies virus neutralizing antibody test after vaccination and the survival rate in mice

The cut-off value of rabies titer, 0.5IU/mL, the minimum defined acceptable threshold as recommended for complete protection against rabies (16). No rabies VNA was detected in placebo group mice. In contrast, vaccinated mice had a small amount of rabies VNA titers after 7 days post a single vaccine injection, and after 14 days, rabies VNA titers of all vaccinated mice reach ≥0.5IU/mL (Fig 3). After challenge with the FB-RABV, all mice had no obvious body weigh change, and all survived, however, in placebo treatment group, mice all developed rabies clinical symptom, and succumbed to death.

**Figure 3.**
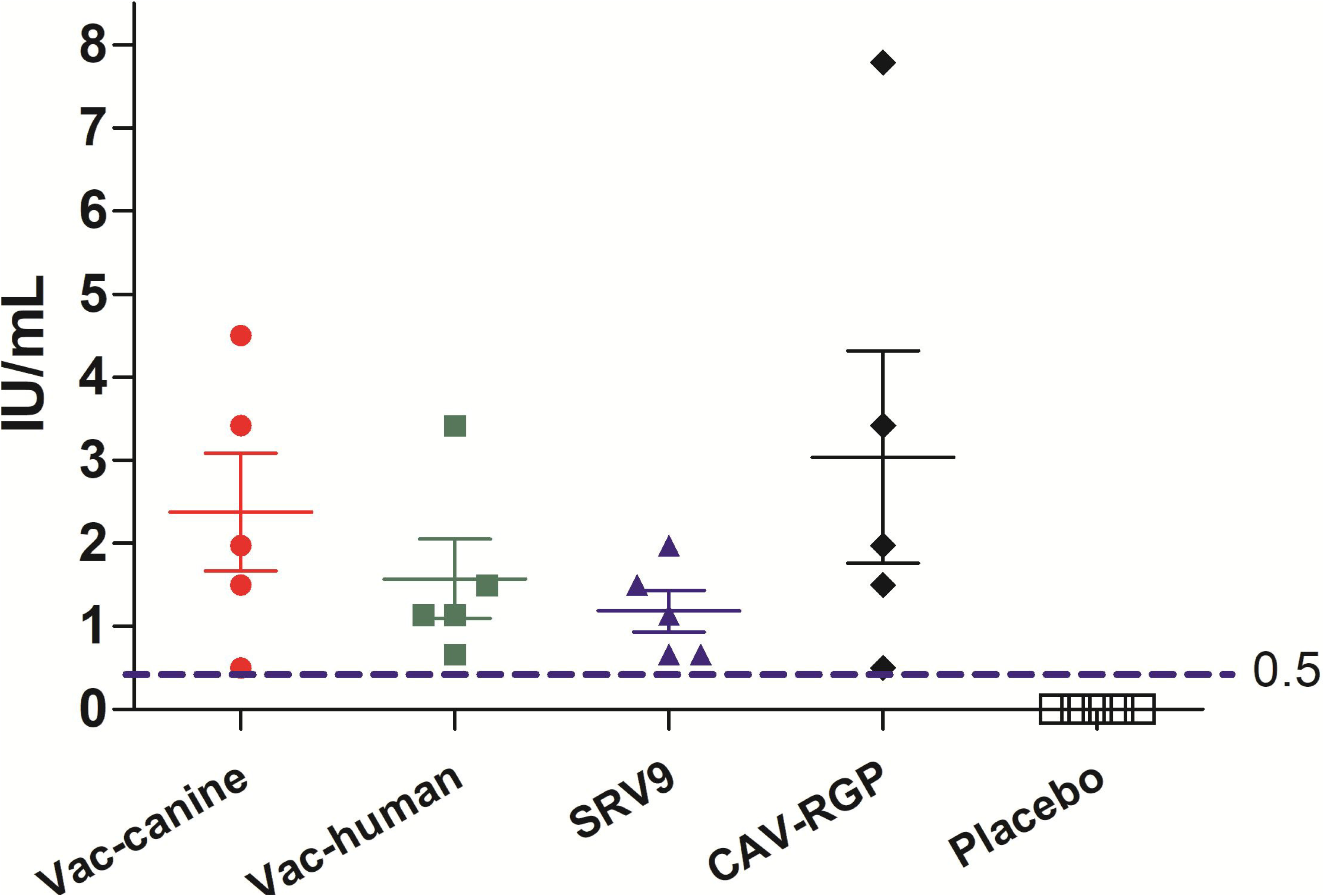
Rabies virus neutralizing antibody (RVNA) titers after 14 days vaccination in mice.

## DISCUSSION

RABV can infect a range of wildlife in specific geographic areas: foxes and raccoon dogs in Europe; foxes in the Middle East; raccoon dogs, camels and ferret badgers in Asia; skunks, foxes, coyotes and mongooses in North America (17); and mongooses in Africa (18–21). Wildlife associated rabies cannot be neglected in a rabies-free country, such as America, where the majority of human rabies are caused by bat bites (22). In Taiwan, an outbreak occurred among ferret badgers during 2012–2013 and a five-decade history of rabies-free humans and domestic animals ended, causing a public health threat (23). In mainland China, where rabid dogs serve as the major source of infection, however, with the increase of vaccination coverage rates to domestic animals and enforcement to the stray dogs, dog-associated human rabies cases rapidly decrease, the wildlife-associated human rabies deaths are now an increasingly recognized problem in southeast China (4), especially that there is no vaccination or any other intervention to prevent FB-related RABV transmission in the animal population, which needs to investigate how RABV was introduced and maintained in the FBs urgently.

Retrospective investigation has revealed that ferret badger-associated rabies was first reported in 1997; however, it first arose in 1994, and it has caused nearly hundreds of human fatalities, though this is likely an underestimated number (24). In this study we focused on the Poyang and Qiandao lake regions where both human and dog rabies co-exists. For the Qiandao lake, FB RABV was only isolated in years 2008, 2012 and 2013. The absence of FB RABV detection from 2009-2011 was possibly due inadequate surveillance. Interestingly, in our 112 FB RABVs positive from the 3,622 samples, 88 were in Lineages A and C: the lineages diverged in 1937 and 1941, respectively. Only 24 isolates were recovered for Lineages B and D, which we estimate as having diverged in 1990 and 1997. We speculate that our better chance (3.7 times) of finding a FB-associated RABV in lineages A and C is because the virus has been circulating for a longer time and is therefore better adapted in the FB population, which might explain, but not proven, its sustained transmission in the Poyang and Qiandao lake regions. Lineages B (year 1990) and D (1997) are corresponding with two major rabies epidemics in China (1980–1990, and 1997– present). Lineages A and C diverged from dog RABVs in the 1930s and 1940s when there was no national disease reporting system. However, TMRCA of China canine RABVs is estimated to be about 1745 CE in our calculation and other studies, suggesting RABV circulation in the FB population occurred after the emergence of dog rabies virus (20, 25).

Numerous studies have shown that various wildlife can be colonized by dog RABV variants to become successful rabies host. Typical examples include the cases in Europe where rabies evolved from domestic dogs to raccoon dogs and foxes (26), the emergence of fox rabies in Colombia and Brazil (27–29), and wildlife rabies in the United States (30). Other spillover events of rabies from dogs to wildlife have been observed in southern Africa (31–34), Puerto Rico (35, 36) and Turkey (37). Because wildlife RABV variants can also transmit to unvaccinated dogs, thus entering urban environments, as has been observed in former Yugoslavia (38). Our study supports the hypothesis that the rabies epizootic in the FB population is a consequence of dog RABV spillover at different times through multiple introductions. Rabies may have been circulating in the FB population for over 60 years (1937) before the first human death was reported in the 1990s. China’s priority for rabies control should focus on dogs, a complicating factor that needs to be considered is rabies in wildlife. It is noteworthy in our study that a canine RABV isolate from Zhejiang province was classified into FB Lineage D, which is composed of FB-RABV isolates (Fig 2). This could represent a cross species transmission event in which either FBs introduced RABV to the dog population or dogs introduced RABV to FBs. Further investigation is needed when more RABVs are isolated from both animal species in the same region.

**Fig 2.**
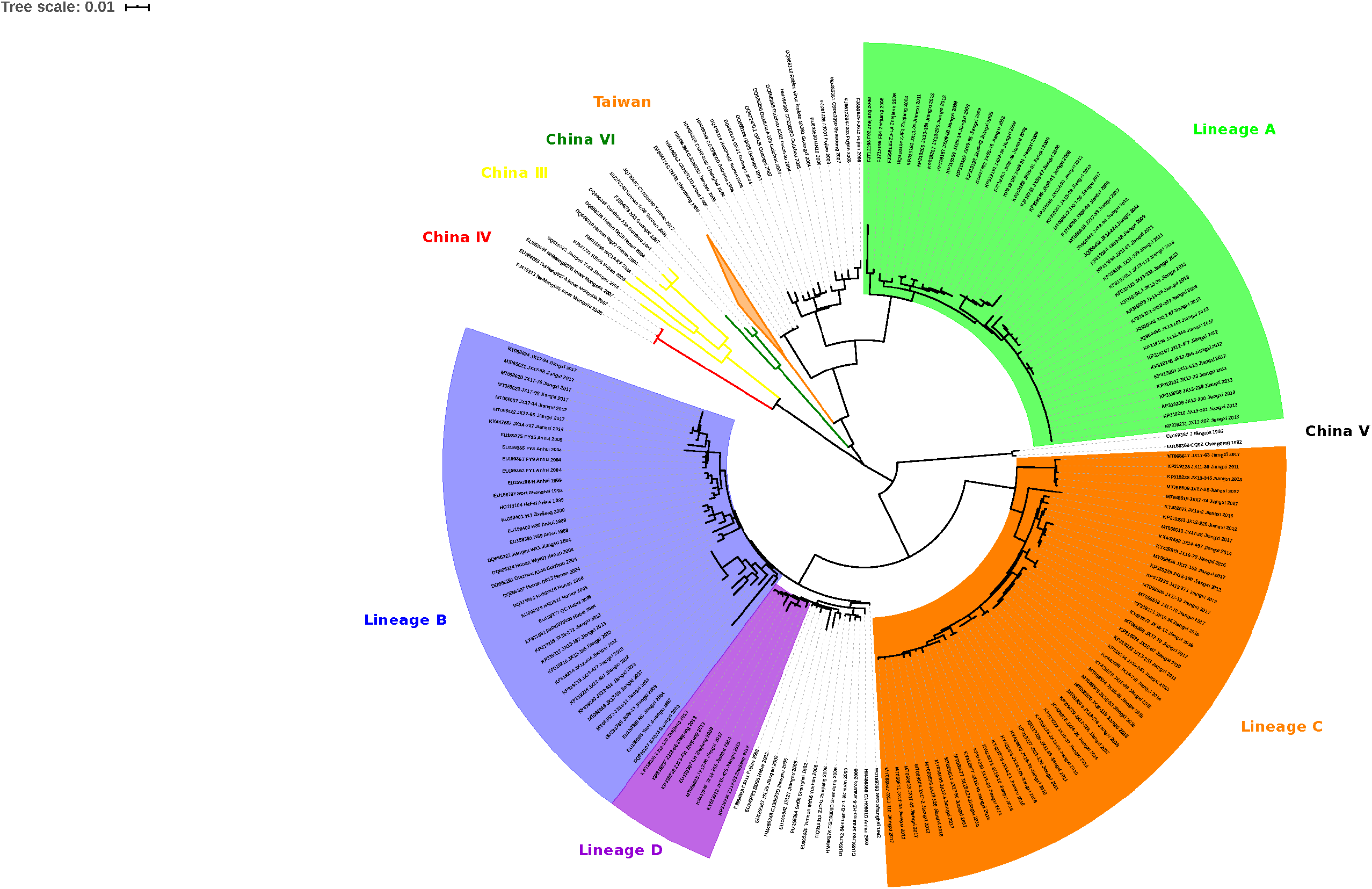
Maximum likelihood phylogeny of 244 RABV sequences separates the 112 ferret badger associated RABVs into 4 distinct Lineages: A, B, C and D. Lineage A was rooted with China dog RABV Group II, Lineages B and D with China dog RABV Group I, and Lineage C was independent, sitting in the middle of Group I and II. The numbers below the branches were bootstrap values (%) of 1,000 replicates.

For rabies virus, selective pressures such as adaptation in a new host species favor the emergence of mutants with viral infectivity and pathogenicity change (39). During the period when rabies virus adapted in ferret badger, our results demonstrated that the pathogenicity of FB-RABV variants decreased to humans and other animals. The laboratory studies were conducted in mice and dogs, mortality-rates ranging from 20% to 100% in mice and 0 to 20% in beagles in the groups given the various isolates, which may indicate that FB-RABV variants can induce inapparent infections in mice and dogs. A previously serological survey of ferret badgers in southeast China found that 69.6% of 57 animals had serum neutralizing antibodies of rabies virus, which may be due to inapparent infections with rabies virus in the FB population (11).

Evidence from laboratory and field studies suggested that inapparent rabies infection may occur in animals (40). It has been suggested that the RABV strain causing enzootic rabies in Ontario in 1950s is a variant with low pathogenicity in humans and dogs (41). No further study to provide an explanation for the phenomenon, so the evidence for inapparent rabies infection in humans was indefinite (40, 42). The explanation is that the persons who were bitten by rabid animals should receive postexposure prophylaxis (PEP) for rabies, thus the inapparent infection were covered up. In this study, the human volunteers of Jiangxi province were selected from the people who frequently contacted with FB and the duration was more than three years. We found nine persons, who participated in the survey, had been bitten by FB and never received rabies vaccines, 5 had detectable rabies neutralizing antibodies with 3 having protective level of antibody titres (≥2.6 IU/ml). This is the first definite evidence that the five human volunteers have been exposed to rabies virus through FB bites, and experienced an inapparent rabies infection without illness. Further studies are needed to proof whether these human subjects had inapparent rabies infection.

Nowadays, dog-associated human rabies cases decline rapidly with sufficient production levels for rabies vaccination of domestic dogs and PEP administrations enhanced, FB-associated rabies becomes to be a major threaten factor in some areas of China(43). Our study demonstrates that only one dose vaccine either inactivated vaccine, attenuated vaccine or recombined submit vaccine can be used to control rabies in ferret badger.

In conclusion, our study in the Poyang and Qiandao lake regions for the last 11 years suggests that RABV can be maintained in the FB populations without human intervention. The TMRCA of FB associated rabies coordinates with canine rabies epidemics in China, and rabies in FBs is a later event after multiple introductions from canine rabies virus. Rabies control in FBs poses a serious immediate concern in China. The potential of FB becoming an animal model, and the viral susceptibility of FB-RABVs to other animals, and mechanism of inapparent infection need further study. Together, Investigation of the phylogenetic and evolution of ferret badger-associated rabies virus strains over the course of an epidemic can help in the development of strategies to combat and control viral disease (44, 45). Further, more research should be devoted to the development of oral vaccines for stray dogs and ferret badgers.

## MATERIALS AND METHODS

### Ethics statement

All animals experiments described in this paper have been conducted according to the Guideline on the Humane Treatment of Laboratory Animals stipulated by the Ministry of Science and Technology of the People’s Republic of China (MOST) (46) and approved by the Animal Welfare Committee of the Institute of Military Veterinary Medicine, Changchun, China. All animals were housed in a climate-controlled laboratory with a 12 h day/12 h night cycle.

Mice were purchased from Changchun Institute of Biological Products, China. Healthy, non-rabies vaccinated, 5-month old healthy female beagles were obtained from Bolong experimental animal Co., LD., Qingdao, China. All dogs were housed individually in temperature- and light-controlled quarters in the Animal Facility, Jilin University.

### Sample collection

From May 2008 through 2018, 3,622 FB brain tissue samples were collected across the Poyang lake (Jiangxi province) and Qiandao lake regions (Zhejiang province) (Figure 1). Most of the samples were obtained from the Poyang lake Region, because this is the most densely inhabited FB area in China. Both dead and living FBs were collected from the field and their heads (after euthanization if still alive) were shipped to our laboratory for rabies detection and RABV isolation.

### RNA extraction, RT-PCR and sequencing

Total RNA of the collected FB brain homogenate was extracted by a single-step method using Trizol Reagent (Invitrogen, Life Technologies, USA) according to the manufacturer’s instructions. For nucleic detection, One-Step-Reverse transcription (RT)-polymerase chain reaction (PCR) was used to obtain the lyssavirus sequences as previously described (47), As for RABV positive template RNA, One-Step-RT-PCR was performed with the designed primer sets according to the nucleoprotein (N) gene sequence of FB-RABV JX08-45 (GenBank: GU647092), primers cover the ORF of N gene: N-F (1-22): 5’-ACGCTTAACAACAAAATCAGAG-3’; N-R (1519-1540): 5’-CTCGGATTTACGAAGATCTTGC-3’.

The PCR conditions used were: reverse reaction at 50°C for 30min; denaturation at 94°C for 2min, then 35 cycles at 94°C for 30s, 52°C for 30s, and extended at 72°C for 1min30s, followed by a final extension at 72°C for 7min. After 1% agarose gel electrophoresis, the expected PCR bands were purified, then the products were sequenced twice using the N-F/R primers (Sangon Biotech Co., Ltd., Shanghai, China).

### Phylogenetic analysis

A data set of 244 complete N gene sequences from China RABVs was compiled (S1 Table). Similarity scores and percentage identities of the sequences were calculated using the software DNAStar. Maximun Likelihood (ML) analysis was performed using MEGA version 8.0 (48). Bootstrap support was estimated using 1,000 replicates.

### Evolutionary analysis

The Bayesian Markov Chain Monte Carlo (MCMC) method in the BEAST v1.7.5 package (http://beast.bio.ed.ac.uk) was used to reconstruct the Maximum Clade Credibility (MCC) phylogenetic tree, estimate the nucleotide substitution rate (per site, per year), and calculate the Time of the Most Recent Common Ancestor (TMRCA) after sequence alignment (49, 50). Mutation rate was determined under a relaxed uncorrelated lognormal molecular clock model using the general time reversible (GTR) substitution model, incorporating gamma distribution to include in the estimations the variation rate among sites (Γ4), and invariable sites (I). The nucleotide substitution model that best fitted the data set was determined by jModelTest (51). All chains were run for 50 million generations to ensure sufficient mixing. Parameters and trees were recorded every 10,000 steps. An effective sample size (ESS) >200 was achieved for all estimated parameters. The Highest Posterior Densities (HPD) were calculated with 10% burn-in and checked for convergence using Tracer (v1.5).

### Virus isolation (Mouse inoculation test, MIT)

FB brain tissues testing positive for rabies by RT-PCR were homogenized as a 10% suspension in Dulbecco’s modified Eagles’s medium (DMEM; Gibco, CA, U.S.) supplemented with antibiotics (100IU/ml penicillin and 100μg/ml streptomycin). The supernatant fluids were clarified by 8,000×g for 10min and aliquots of 25μl were injected intracerebrally into 3-day-old suckling mice (Balb/c mice, Changchun Institute of Biological Products, China) according to an approved animal protocol (52). Animals were observed daily for 21 days post-injection and those showing symptoms of rabies were euthanized. Brain tissues were removed and tested for RABV antigen by the direct fluorescent antibody (DFA).

### Pathogenicity of FB-RABV isolates in mice and beagles

Eight FB-RABV strains used in this experiment (JX08-45, JX13-189, JX09-17, JX13-417, JX10-37, JX13-228, ZJ12-03, ZJ13-431) were isolated in Jiangxi (Jingdezhen, Fuzhou and Xiushui) and Zhejiang (Taizhou) province. RABV BD06 (GenBank accession no. EU549783.1) was isolated in our laboratory in 2006 from a rabid dog in China. This virus is maintained in dogs and it was shown in our previous experiments to cause 80% mortality in unvaccinated dogs after intramuscular injection. In addition, this virus strain is responsible for the majority of rabies cases among humans and dogs in China (53). All FB-RABV isolates were from brain tissues and were passaged three times intracerebrally in 3-day-old suckling mice (Balb/c mice, Changchun Institute of Biological Products, China). When moribund, the mice were euthanized. 10% (w/v) suspensions were homogenized in DMEM containing 10% newborn calf serum and antibiotics, centrifuged to remove debris, and the supernatants were stored at −80°C.

Titers of the viral isolates were determined by intracerebral (IC) injection of 10-fold dilutions into 5-week-old female Balb/c mice, and median mouse IC lethal doses (MICLD_50_) were calculated using the Reed-Muench formula. Five 5-week-old female Balb/c mice per group were intramuscularly inoculated with 0.1 ml of 10^4^ MICLD_50_ of each strain into the left thigh muscle. Control animals were mock infected with 0.1 ml of DMEM containing 10% calf serum and antibiotics. The mice were observed twice daily for 28 days post inoculation (dpi): early morning, when body weights were measured, and late afternoon. Incubation period was taken as the number of days from injection to first appearance of neurological symptoms such as tremors or paralysis of one hind leg. In cases when death was imminent in the late afternoon, animals were euthanized and the morbidity period was taken as being the next day.

For further evaluate the pathogenicity in dogs, four groups (JX13-189, JX13-228, ZJ13-431, BD06) of female beagles (n = 5) were inoculated intramuscularly (IM) into both left and right masseter muscles. Each inoculated animal received a total of 10^4.4^ MICLD_50_ of each viral isolate. Control animals were injected with DMEM containing 10% calf serum and antibiotics. Ninety days postinoculation (PI) all surviving animals were challenged with rabies virus (canine street virus; 1×10^5.4^ MICLD_50_) by the inoculation of 0.5ml each in the left and right masseter muscles. The brainstem was collected at necropsy and tested by DFA. After challenge, the beagles were observed for another 90 days, and all surviving animals were euthanized to collect brain samples and tested by DFA.

### Serological survey among humans who exposure to ferret badger RABV

Nine sera were obtained from the volunteers collected during an investigation of humans bitten by FB more than twice during the three years and with no rabies PEP treatment. All participants involved in the study had written informed consent. Sera were separated by centrifugation following incubation at ambient temperature for 3h, inactivated at 56°C for 30 min, each sample was tested for rabies virus-specific neutralizing antibodies (VNA) by the standard fluorescent antibody virus neutralization (FAVN) test using BHK-21 cells (16).

### Rabies vaccination in mice and challenge with ferret badger rabies virus

To determine the availability of emergency prophylaxis to the exposure to FB rabies, 25 Balb/c mice were randomly divided into 5 groups and immunized intramuscularly with a single injection 100μl (containing NIH potency 0.2IU) of canine inactivated vaccine produced by Jilin Heyuan Bioengineering Co. Ltd, human inactivated vaccine produced by Liaoning Chengda Bioengineering Co. Ltd, and other vaccination produced by our lab (attenuated SRV9 and a recombinant Canine Adenovirus Vaccine CAV-2-E3D-RGP), the placebo group was injected with DMEM as control. After 14 days post vaccination, mice were challenged with 0.1 ml of 10^4^ MICLD_50_ of FB-JX08-45 into the left thigh muscle, and housed for another 28 days, all survival mice were sacrificed after serum collection. Sera were separated and rabies virus neutralizing antibody (RVNA) was assayed using the standard FAVN method at 7 and 14 days post vaccination.

### Accession numbers

The new gene sequences in this study were submitted to NCBI GenBank nucleotide sequence database, accession numbers supplied as S1 Table.

## SUPPLEMENTAL MATERIAL

Supplemental material is available online only.

SUPPLEMENTAL FILE 1, PDF file, 0.35 MB.

## ACKNOWLEDGMENTS

We thank Pro. Xianfu Wu, Centers for Disease Control and Prevention, Atlanta, America, for editing and revising the manuscript.

This work was funded by the National Natural Science Foundation of China (31802202 and 31941016) and the National Key Research and Development Program of China (Grant No. 2016YFD0500401). The funders had no role in study design, data collection and analysis, decision to publish, or preparation of the manuscript.

